# Prediction of soil probiotics based on foundation model representation enhancement and stacked aggregation classifier

**DOI:** 10.1101/2025.06.13.659431

**Authors:** Qiang Kang, Haotong Sun, Yayu Wang, Xiaolong Fang, Yuxiang Li, Yong Zhang, Tong Wei, Peng Yin

## Abstract

Soil probiotics are indispensable in agro-ecosystem, which enhances crop yield through nutrient solubilization, pathogen suppression, and soil structure improvement. However, reliable prediction methods for soil probiotics remain absent. In this study, we utilize genomic foundation models to generate representations from samples’ sequences, and then, enhance them by deeply integrating domain-specific engineered features. The enhanced representations enable training a powerful classifier for a target task, instead of common parameter fine-tuning. Inspired by the stacking ensemble learning framework, we also design a stacked aggregation classifier. It predicts the label of a sample with only leveraging partial sequence segments from this sample, effectively addressing the challenges in processing long sequences. The proposed method is applied on prediction of soil probiotics and obtains 97.50% accuracy and 0.9807 AUC value on balanced and imbalanced test sets, respectively. Potential functional genes are revealed from the predicted probiotics, providing biologically insights for more related studies.

## 1. Introduction

Soil probiotics, encompassing functionally active microbial consortia such as *Bacillus* and *Pseudomonas*, serve as indispensable regulators of agro-ecosystems [1,2]. These microorganisms can drive biogeochemical cycles and enhance crop productivity through multifunctional mechanisms. *Azotobacter* has been reported to fix approximately 20 kg N/ha/per year, contributing significantly to soil nitrogen content [3], while *Pseudomonas fluorescens* can produce 2,4-diacetylphloroglucinol to suppress *Ralstonia solanacearum* and has the ability to solubilize phosphate [4]. In field applications, inoculating cold-tolerant phosphate-solubilizing *Pseudomonas* strains increased wheat yield by approximately 22% under natural growing conditions [5]. Targeted discovery and cultivation of soil probiotics is critical for sustainable agriculture, enabling innovations such as microbial consortia-based pesticides and fertilizers that synergistically improve soil resilience and crop health [6–8].

Traditional methods for identifying probiotics rely on culturing bacteria in selective medium and analyzing their 16S rRNA gene sequences [9–12]. However, these methods cannot fully reveal the functional capabilities of probiotics, especially since only a small fraction of soil microorganisms can be grown in laboratory conditions [13,14]. In recent years, artificial intelligence (AI) has made significant strides life sciences, with various models being widely applied to species classification and identification tasks [15–17]. Compared to traditional methods, AI-driven methods can learn deep information from microbial genomic sequences to identify candidate strains with desired functional traits, significantly reducing time and cost.

For prediction of probiotics, methods such as iProbiotics and metaProbiotics have been developed. IProbiotics is a machine learning-based tool that utilizes k-mer features and incremental feature selection to predict potential probiotic strains from genomic data, achieving high accuracy [18]. MetaProbiotics employs natural language processing techniques to represent DNA sequences as word vectors and uses random forest classifiers to predict probiotic-associated bins from metagenomic data [19]. However, these methods train the models using data from specific environments, such as the human gut. Due to significant differences in microbial communities across environments (for example, soil microbiomes exhibit greater complexity and diversity compared to human gut microbiomes [20]), these methods may not be applicable to soil microorganisms. Furthermore, retraining these models by directly replacing the training data with soil microorganism may present unreliabilities due to their original designs inherently coordinate sample scale requirements, data preprocessing protocols, and sample representation strategies for a specific task. Currently, there is a lack of AI-driven methods specifically designed for predicting soil probiotics, and developing such a method is necessary and meaningful.

Large language models represented by BERT [21] and the GPT series [22–25] have rapidly advanced in natural language processing. Their architectures have subsequently been adapted to decipher the genomic “language”, thereby establishing several foundation models. One of the pioneering models is DNABERT, which applies the BERT architecture to DNA sequences using k-mer tokenization [26]. DNABERT-2 introduces improvements such as replacing k-mer tokenization with Byte Pair Encoding and incorporating Attention with Linear Biases [27]. Evo comprises 7 billion parameters and is trained on 2.7 million prokaryotic and phage genomes [28]. Its enhanced version, Evo2, expands the parameters to a maximum 40 billionis and is trained on 9.3 trillion DNA base pairs from a highly curated genomic atlas spanning all domains of life [29]. The Nucleotide Transformer represents a family of transformer-based models developed on integrated information from 3,202 human genomes and 850 genomes from diverse species, with parameter sizes ranging from 50 million to 2.5 billion [30]. Additional models such as HyenaDNA [31], GenomeBert [32] and FGeneBERT [33] demonstrate diversified applications, collectively underscoring the significant contributions of foundation models in genomics research.

The application of genomic foundation models to target tasks such as prediction of probiotics typically relies on fine-tuning [21]. However, fine-tuning demands large-scale and high-quality data, and specialized expertise in architecture design and hyperparameter optimization, coupled with access to high-performance computing clusters (e.g., multi-GPU systems with NVLink interconnects) [34], which creates barriers for users lacking professional support, such as biologists and doctors. A critical technical limitation of fine-tuning arises from catastrophic forgetting [35], where models rapidly discard previously learned information when adapting to new tasks. It reduces model reliability in multi-task applications. While low-rank adaptation [36] shows promise in preserving core knowledge, its implementation further compounds the technical complexity. Whole-genome sequences of microorganisms exceed the context window limitations of most Transformer-based architectures [37]. While Hyena-based architectures mitigate these limitations through implicitly parametrized long convolutions and data-controlled gating [31], they still fail to cover the whole-genomic sequences of many microorganisms. Furthermore, de novo genome assembly pipelines are prone to errors due to repetitive regions and sequencing biases, which may introduce errors prior to model training [38].

To address these challenges, we present a method based on foundation model representation enhancement and stacked aggregation classifier and apply it for prediction of soil probiotics. We leverage the foundation models for inference only on the target task, followed by enhancement of their output representations by deeply integrating domain-specific engineered features as an alternative to fine-tuning. It enables to train a powerful classifier for a target task, significantly reduces computational resource requirements and time consumption, while being easy to implement and more user-friendly. Furthermore, we divide long sequences into segments and focus on stacking enhanced representations of partial segments to represent the entire sample. It not only circumvents issues arising from processing length sequences and assembly errors but also enables reliable predictions even for samples with incomplete sequencing coverage. We conduct extensive experiments based on diverse foundation models with different parameter sizes and architectures to systematically evaluate the feasibility and effectiveness of the proposed method. We further explore potential functional genes and their enriched pathways from the predicted probiotics, which provides actionable insights for subsequent wet-lab experiments.

## 2. Methods

The overview of the proposed method is shown in Figure 1. Taking a bacterial sample as an example, its genomic sequence is divided into segments, with partial segments selected. These selected segments are input into a pre-trained foundation model to generate representations, while engineered features are extracted from these segments. The vectors of foundation model representation and engineered feature are aligned, after which the engineered features are deeply integrated into the foundation model representations for enhancement. The stacked aggregation classifier comprises two sub-classifiers. The first-level classifier processes each segment’s enhanced representations to output a score, and the scores from all segments are aggregated into a vector. This score vector is then input into the second-level classifier, which output the final label and score of the sample.

**Figure 1.**
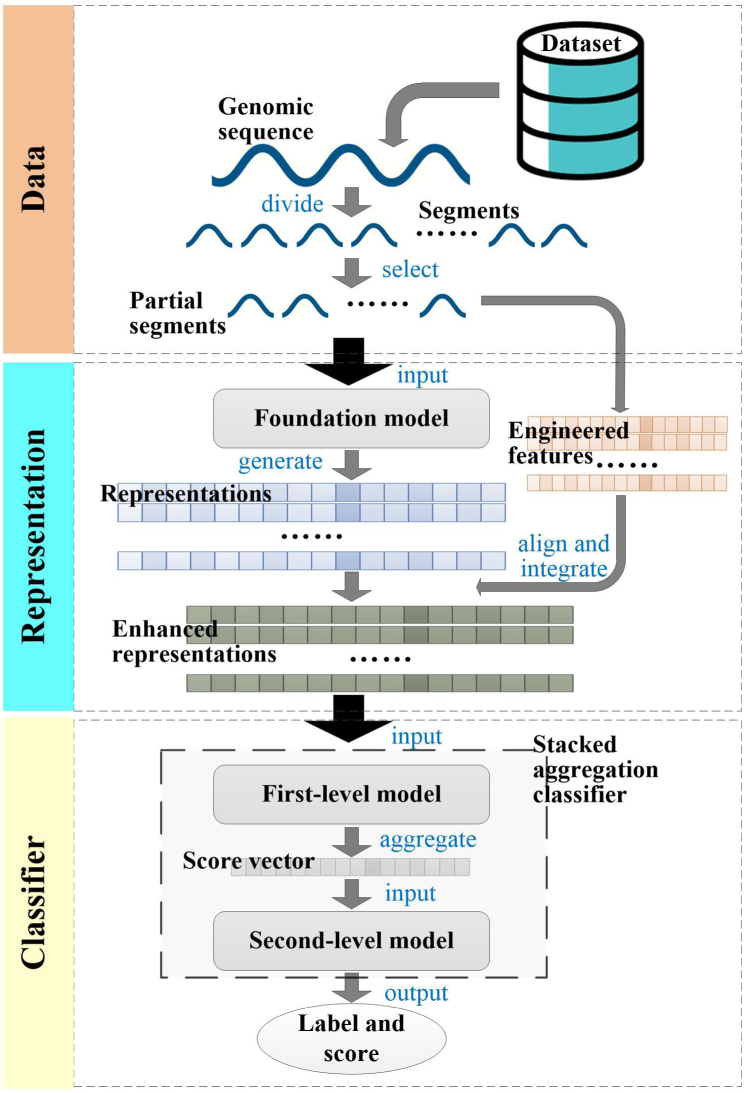
Overview of the proposed method. The genomic sequence of a bacterial sample is divided into segments. Its partial segments are input into a pre-trained foundation model to generate representations, and engineered features are extracted from these segments. The foundation model representation and engineered feature vectors are aligned, and then, the foundation model representations are enhanced by deeply integrating the engineered features. The enhanced representations are fed into the stacked aggregation classifier. The first-level classifier processes each enhanced representation to obtain a score. All scores are aggregated into a vector, which is input into the second-level classifier to output the final label and score.

### 2.1 Datasets and preprocessing

All genomic sequences are downloaded from National Center for Biotechnology Information (NCBI). Positive samples consist of reported soil probiotics, including diverse species such as *Bacillus subtilis*, *Bacillus amyloliquefaciens*, and *Paenibacillus peoriae*. Negative samples are reported soil pathogens. To address the limited number of negative samples, we expand them by: 1) downloading bacterial genomes of the same species as these pathogens from different environments, 2) randomly selecting 100 samples per species (or all available samples if <100), and 3) retaining samples with 95-99% Average Nucleotide Identity to any pathogenic strain via fastANI analysis [39]. For quality control, samples with sequence lengths <1MBP or >10Mbp are filtered out. Each sequence is divided into 1,000bp segments with 200bp overlap between adjacent segments, and the final segment is filtered out if its length <1,000 bp.

All samples are divided into four datasets (a training set, a training-validation set, a balanced test set, and an imbalanced test set), where the training set and training-validation set are employed for trainings in distinct modules, with the training-validation set also utilized for cross-validation. The imbalanced test set simulates real soil environment where probiotics constitute a minority. The details of datasets are shown in Table 1.

**Table 1.** Details of datasets.

| Dataset | #Pos | #Seg_Pos | #Neg | #Seg_Neg |
| --- | --- | --- | --- | --- |
| Training set | 200 | 1,284,328 | 200 | 1,379,878 |
| Training-validation set | 120 | 726,088 | 120 | 839,687 |
| Balanced test set | 80 | 518,285 | 80 | 557,708 |
| Imbalanced test set | 18 | 115,293 | 180 | 1,265,553 |
| #Pos: number of positive samples, #Neg: number of negative samples, #Seg_Pos: number of segments of positive samples, #Seg_Neg: number of segments of negative samples. |  |  |  |  |

### 2.2 Foundation model representation generation

Three cutting-edge genomic foundation models with different parameter sizes and architectures, Nucleotide Transformer-50M [30], DNABERT-2-117M [27] and EVO-7B [28], are utilized to generate diverse representations. A biological sequence is tokenized into discrete subunits and processed by a foundation model’s encoder to generate contextual embeddings. These embeddings form a matrix where each row corresponds to a token’s feature vector. The embedding of the entire sequence is derived from the first positional token, which integrates global sequence information through the attention mechanism, serving as the generated representation. Given an input sequence segment *Seq*=*n*^1^*n*^2^…*n*^1000^ (where *n^i^*∈{*A*,*T*,*C*,*G*} denotes the *i*-th nucleotide), Nucleotide Transformer-50M generates the 512-dimensional representation vector as ***R****_nt_*=[*r* ^1^,*r* ^2^,…,*r*_nt_ ^512^], DNABERT-2-117M produces the 768-dimensional representation vector as ***R****_db_*=[*r_db_*^1^,*r_db_*^2^,…,*r_db_*^768^], and EVO-7B constructs the 4096-dimensional representation vector as ***R****_evo_*=[*r_evo_*^1^,*r_evo_*^2^,…,*r_evo_*^4096^], where *r_nt_^i^*, *r_db_^j^* and *r_evo_^k^* denote the representation values at the *i*-th, *j*-th and *k*-th dimensions of ***R****_nt_*, ***R****_db_*, and ***R****_evo_*, respectively. For each sample, the corresponding three foundation model representations can be expressed as:

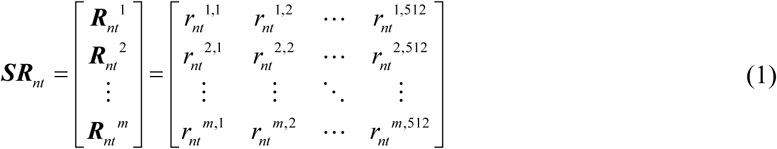

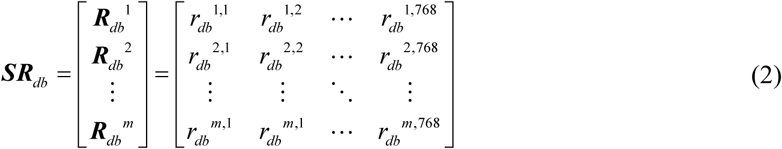

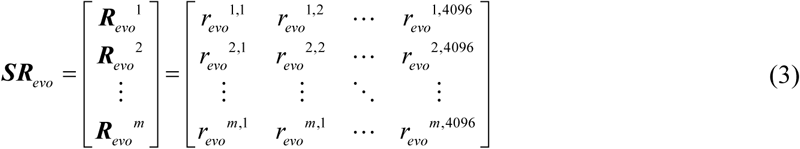

where ***R_nt_****^l^*, ***R****_dp_^l^* and ***R****_evo_^l^*denote different representations of the *l*-th segment and *m* denotes the number of segments.

### 2.3 Engineered feature extraction

Domain-specific engineered features, k-mer and g-gap, are extracted from each sequence segment. K-mer represents the frequency of consecutive *k* nucleotides (e.g., a 3-mer category can be AGC), while g-gap denotes the frequency of discontinuous nucleotides separated by *g* gaps (e.g., a 2-gap category can be A**C, where “*” represents any nucleotide) [18,40]. Consequently, each k-mer category corresponds 4*^k^* features, and each g-gap category matches 4^2^ features. A sliding window is applied to scan each sequence segment, recording the occurrence counts of each k-mer category and g-gap category. These counts are normalized to generate corresponding feature values as:

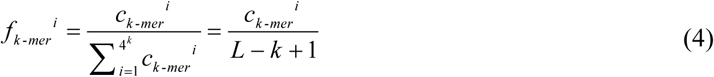

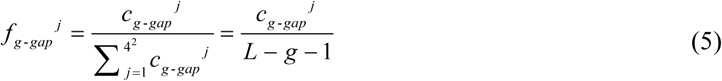

where *c_k-mer_^i^* denotes the counts of the *i*-th k-mer category, *c_g-gap_^j^* denotes the counts of the *j*-th g-gap category, and *L* denotes the sequence segment length. Given a sequence segment, 1, 2 and 3-mers features and 1, 2 and 3-gaps features are extracted to form a 132-dimensional vector ***F***=[*f*^1^,*f*^2^,…,*f*^132^]=[*f_k-mer_*^1^,*f_k-mer_*^2^,…,*f_k-mer_*^84^,*f_g-gap_*^1^,*f_g-gap_*^2^,…,*f_g-gap_*^48^]. For each sample, the corresponding engineered features can be expressed as:

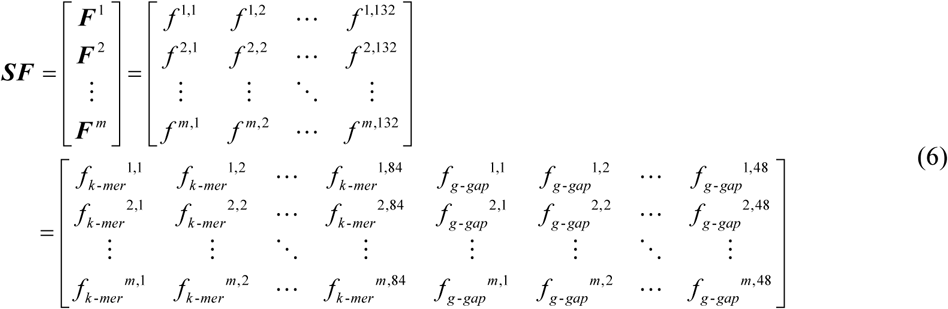

where ***F****^l^*denotes the engineered feature vector of the *l*-th segment.

### 2.4 Foundation model representation enhancement

To enhance the foundation model representations by integrating engineered features, they need to be aligned in dimension and scale. For each sample’s ***SF***, global Min-Max Normalization is applied to standardize its scale as:

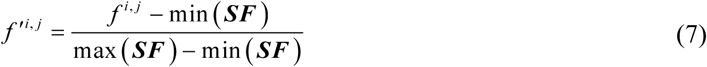

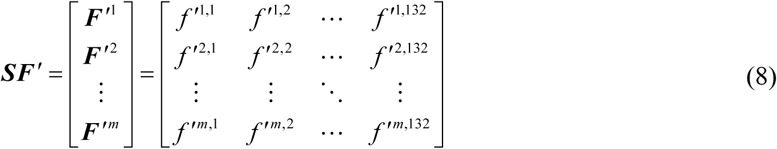

where *f^i,j^* denotes the feature value at the *i*-th row and *j*-th column in ***SF*** and *f’^i,j^* is its normalized value, min() and max() denote the minimum and maximum functions, respectively. For each sample’s ***SR****_nt_*, ***SR****_db_* and ***SR****_evo_*, the following process is performed: 1) their value ranges are unified through global Min-Max Normalization similar as Eqs.(7) and (8); 2) each representation vector (i.e., each row) in them is reduced to 132 dimensions via kernel Principal Component Analysis (PCA) [41], ensuring consistency with the dimension of each feature vector in ***SF****’*; 3) global Min-Max Normalization is performed on them again to ensure scale consistency with ***SF****’*. Such the processed representations can be obtained as:

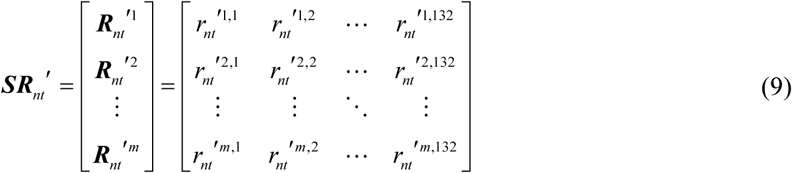

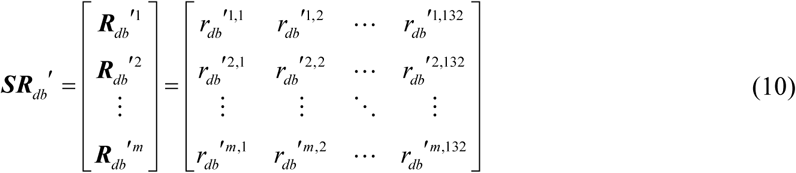

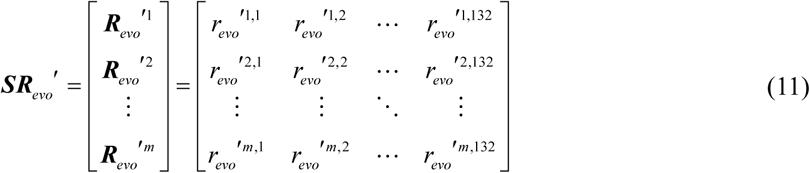

Foundation model representations and engineered features exist in mismatched dimensional spaces, requiring nonlinear integration methods. The cross-attention mechanism can dynamically bridge heterogeneous representations/features through Query, Key and Value interactions [42]. Although cross-attention mechanism introduces an additional training step compared to simple integration methods, the number of trainable parameters it required is far fewer than fine-tuning a foundation model. For the *l*-th segment, its feature vector is integrated into three representation vectors respectively based on multi-head cross-attention mechanism. Here taking ***SR****_nt_’* as an example and the process is similar for ***SR****_db_’* and ***SR****_evo_’*. For each head, Query is derived from the foundation model representation ***R****_nt_’^l^*and Key/Value is derived from the engineered feature ***F****’^l^*, the cross attention can be expressed as:

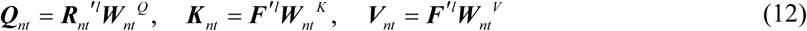

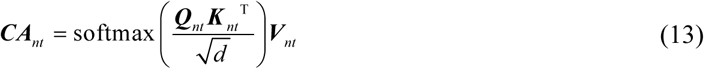

where ***W****_nt_^Q^*, ***W****_nt_^K^* and ***W****_nt_^V^*∈ℝ^132×^*^d^* are the trainable weight matrix with *d*=132/*h*, *h* is the number of heads, the output ***CA****_nt_* is a *d*-dimensional vector. The outputs of all heads are concatenated and linearly projected by a trainable matrix ***W****_nt_^O^*∈ℝ^132×132^, then combined with the original input via residual connection as:

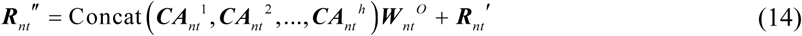

where ***CA****_nt_^i^* denotes the output vector from the *i*-th head, and ***R****_nt_’’* is the enhanced representation vector with 132 dimensions. For each sample, the corresponding enhanced representation can be expressed as:

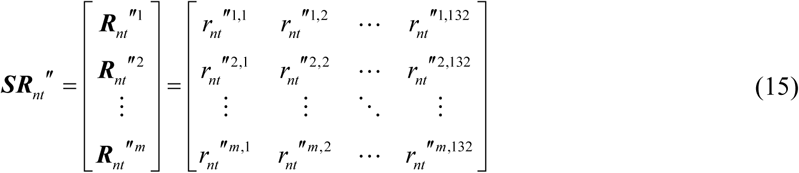

where ***R****_nt_’’^l^*denotes the enhanced representation of the *l*-th segment.

### 2.5 Stacked aggregation classifier

Each sample is represented by an *m*×132 matrix that requires conversion into a vector before being fed into a classifier. The stacking ensemble learning framework integrates the outputs of multiple base learners to generate an ensemble vector, and feeds it into a meta-learner for final output [43]. Inspired by this, a stacked aggregation classifier is designed, which consists of two sub-classifiers. A fixed number of segments are selected from each sample. Each segment’s enhanced representation vector is input into the first-level classifier (similar to the base learner) to output a corresponding score. All scores are aggregated into a vector as input of the second-level classifier (similar to the meta-learner) and the final label and score are output.

The eXtreme Gradient Boosting (XGBoost) model is employed as the first-level classifier. Its gradient-boosted tree architecture recursively learns representation interactions, effectively modeling nonlinear dependencies across sequence segments. The second-level classifier is the Logistic Regression (LR) model. It utilizes L1/L2 regularization to stabilize predictions while maintaining computational efficiency. This stacked aggregation classifier not only achieves a sample representation by partial sequence segments but also has the potential to obtain high accuracy and robustness.

### 2.6 Cascade training strategy and inference process

The proposed method involves trainings in two modules, i.e., cross-attention mechanism and stacked aggregation classifier. The former aims to establish correlations between foundation model representations and engineered features, while the latter needs to build decision boundaries on the enhanced representations. Given their distinct training objectives, they are trained separately with different training sets. Compared to joint training, separate training is easier to implement with less training time, and allows independent module extension. The cross-attention mechanism cannot be trained independently, thus it is followed by a Multilayer Perceptron (MLP) to assist the training, and the loss function is designed as:

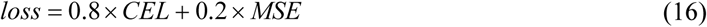

where *CEL* is the cross entropy loss, and *MSE* is the mean squared error for constraining the reconstruction of engineered features. The reconstruction term forces the training to focus on engineered features, preventing over-reliance on foundation model representations. For the stacked aggregation classifier, XGBoost and LR employ their default loss functions.

We adopt a cascade training strategy, which means that the models are trained sequentially, where earlier trained model helps the training of subsequent ones (Figure 2). Each sequence segment inherits the label from its source sample. The cross-attention mechanism is trained with the training set. The aligned foundation model representations and engineered features of all samples’ segments are used (Figure 2A). The stacked aggregation classifier is trained with the training-validation set. We select *m*×5 segments from each sample (*m* denotes the number of segments selected per sample during inference, here expands it to five times to augment the training data), and obtain their enhanced representations based on the trained cross-attention mechanism to train the XGBoost model (Figure 2B). Subsequently, *m* segments are selected five times with replacement from each sample, ensuring the selected segments are not entirely identical at different times and are completely inconsistency with those for training the XGBoost model. It can also augment the training data when the number of training samples is limited. The enhanced representations (obtained based on trained cross-attention mechanism) of the five *m*-segment groups are separately fed into the trained XGBoost model, generating five corresponding *m*-dimensional score vectors to train the LR model (Figure 2C).

**Figure 2.**
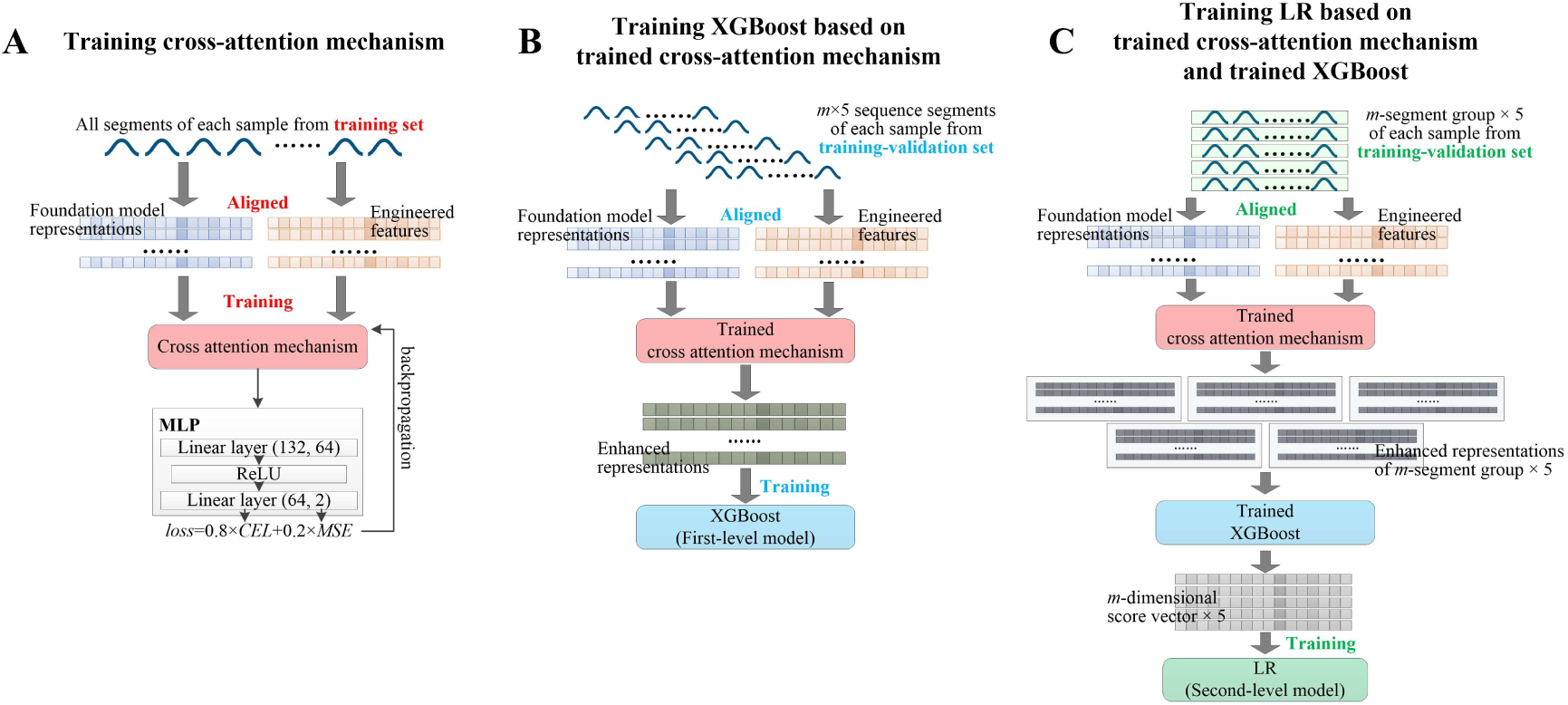
Cascade training strategy. The models are trained sequentially, where earlier trained model helps the training of subsequent ones. (**A**) The cross-attention mechanism is followed by a MLP to assist the training. It is trained using the aligned foundation model representations and engineered features of all sequence segments of each sample from training set. (**B**) For each sample from training-validation set, *m*×5 sequence segments are selected. Their enhanced representations are obtained based on the trained cross-attention mechanism to train the XGBoost model. (**C**) For each sample from training-validation set, *m* sequence segments are selected five times with replacement, and the selected segments are not entirely identical at different times and are completely inconsistency with those for training the XGBoost model. Their corresponding five groups of enhanced representations are obtained based on the trained cross-attention mechanism. These enhanced representations are fed into the trained XGBoost model to generate the five *m*-dimensional score vectors to train the LR model.

In the inference process, only *m* sequence segments from the target sample are selected for using the proposed method to obtain: 1) the foundation model representations, 2) the extracted engineered features, 3) the enhanced representations obtained based on the cross-attention mechanism, and 4) the predicted label and score from the stacked aggregation classifier. For long-sequence samples with abundant segments, multiple *m*-segment groups can be selected for multiple independent predictions, and their results can be considered comprehensively.

### 2.7 Gene function annotation and enrichment analysis

For a target sample, its genes are obtained using Prodigal v2.6.3 [44]. These genes are annotated using KofamScan v1.3.0 [45] (configured with KEGG release 114) to obtain their KO identifiers. These identifiers are filtered according to: 1) null threshold, 2) score<threshold, and 3) *e*-value>1e-5. The filtered identifiers are mapped to functions from the KEGG ORTHOLOGY database [46]. These mappings are cross-referenced with reported phenotypes of the target sample from public literature, which may lead to inference of which genes in the target sample exert critical impacts on specific functional traits. Enrichment analysis is performed using ReporterScore v0.1.9 [47] (configured with KEGG release 114) to identify pathways significantly enriched on these potential functional genes.

### 2.8 Evaluation criteria

True Positive (TP) represents the number of correct predictions for probiotics, False Negative (FN) represents the number of incorrect predictions for probiotics, False Positive (FP) represents the number of incorrect predictions for non-probiotics, and True Negative (TN) represents the number of correct predictions for non-probiotics. The evaluation criteria used for cross-validation and the experiments on balanced test set include Recall (REC), Precision (PRE), Specificity (SPE), Accuracy (ACC), F1-score (F1_S), Matthews Correlation Coefficient (MCC), and Area Under Curve (AUC) from Receiver Operating Characteristic (ROC) curve. For experiments on imbalanced test set, the evaluation criteria include REC, SPE, Geometric mean of Recall and Specificity (GRS), and AUC. The related equations are expressed as:

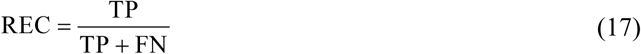

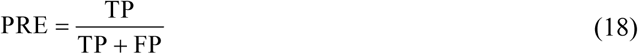

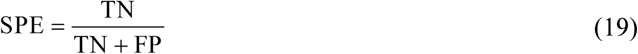

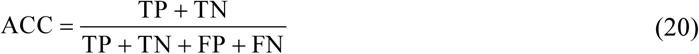

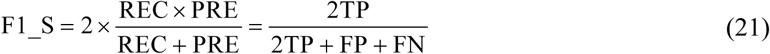

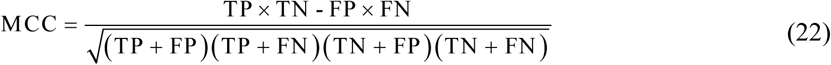

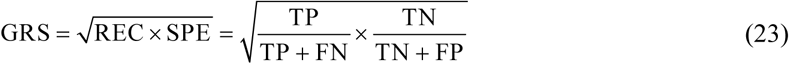

### 2.9 Project implementation

Main scripts are written in a Python v3.8.15 virtual environment on a computing node equipped with four NVIDIA A100 40GB GPUs and 256GB RAM. Prodigal (implemented in C) is compiled with GNU Compiler Collection v14.2.0. KofamScan utilizes Ruby v3.2.2 as its scripting language. ReporterScore is implemented using R v4.4.0.

The k-mer size and segment length of fastANI are set to 16 and 1000, respectively. The number of heads in multi-head cross-attention mechanism is set to 11. In MLP, the first linear layer maps the input from 132 dimensions to 64 dimensions, followed by the ReLU activation function, and the second linear layer reduces the 64-dimensional input to a 2-dimensional output. Prodigal, KofamScan and ReporterScore perform only input and output operations without any adjustments. The hyperparameter settings of XGBoost and LR are described in the section 3.3.

## 3. Results

### 3.1 Investigation on number of segments for representing a sample

From the target sample, *m* segments are selected as the input. If *m* is too small, the sample may be represented inadequately, and if *m* is too large, resource consumption increases. We freeze the parameters of the cross-attention mechanism and conduct 5-fold cross-validation on the training-validation set (in each fold, 10 independent experiments are performed, and each experiment selecting *m* segments that are not completely consistent with previous ones to obtain statistical results) to observe how varying *m* affects the prediction (Figure 3).

**Figure 3.**
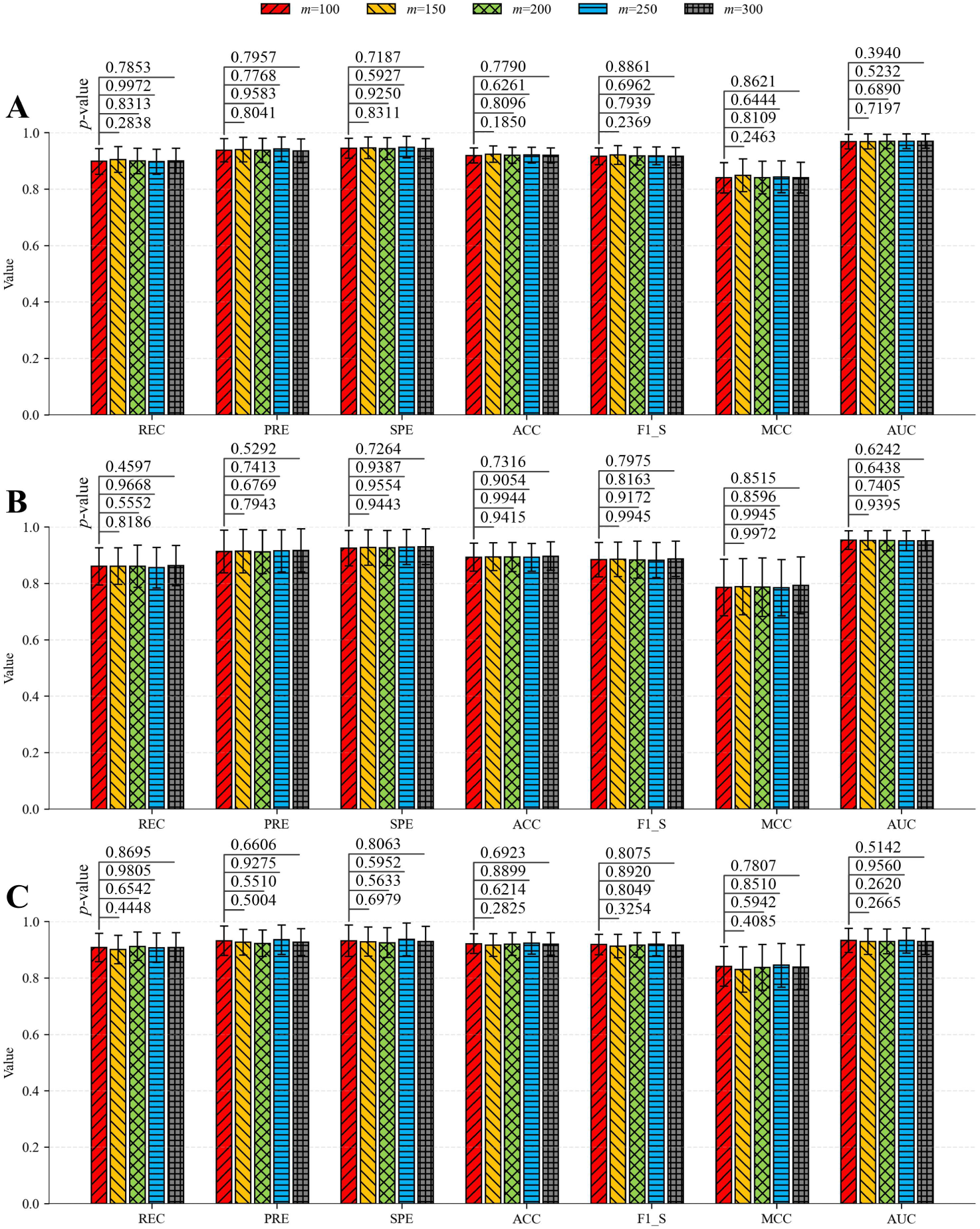
Effect of the number of selected segments (*m*) on prediction. The values above the bars are *p*-values from Mann-Whitney U tests comparing results at *m*=100 versus other *m*-values. (**A**) The foundation model representations are generated by Nucleotide Transformer-50M. (**B**) The foundation model representations are generated by DNABERT-2-117M. (**C**) The foundation model representations are generated by EVO-7B.

When *m* takes different values, the proposed method shows minimal variations in each evaluation criterion, regardless of which foundation model generates the representations. Statistical tests reveal no significant difference (*p*-value>0.05) between results at *m*=100 and each of other *m*-values. We therefore set the number of segments selected from a target sample to 100. This achieves a moderate reduction in computational resource and time consumption while maintaining applicability, and even for an incomplete sample sequence, prediction can be executed as long as 100 segments can be partitioned.

### 3.2 Effectiveness evaluation of enhanced representation and stacked aggregation classifier

The inputs to the stacked aggregation classifier can be either the enhanced representation vectors of samples, the original engineered feature vectors, or the original foundation model representation vectors. For the enhanced representation matrix of a sample, in addition to using stacked aggregation to convert it into a vector as classifier input, approaches such as taking the mean, maximum, or minimum along each dimension of the matrix, or applying PCA can also be employed. Furthermore, a classifier can be trained using segments’ vectors of the training samples, and then, the trained classifier is used to predict labels for each segment of the target sample, with voting ultimately determining the final label. We adopt the same 5-fold cross-validation scheme as in the section 3.1 to evaluate the effectiveness of enhanced representations and stacked aggregation classifier (Figure 4).

**Figure 4.**
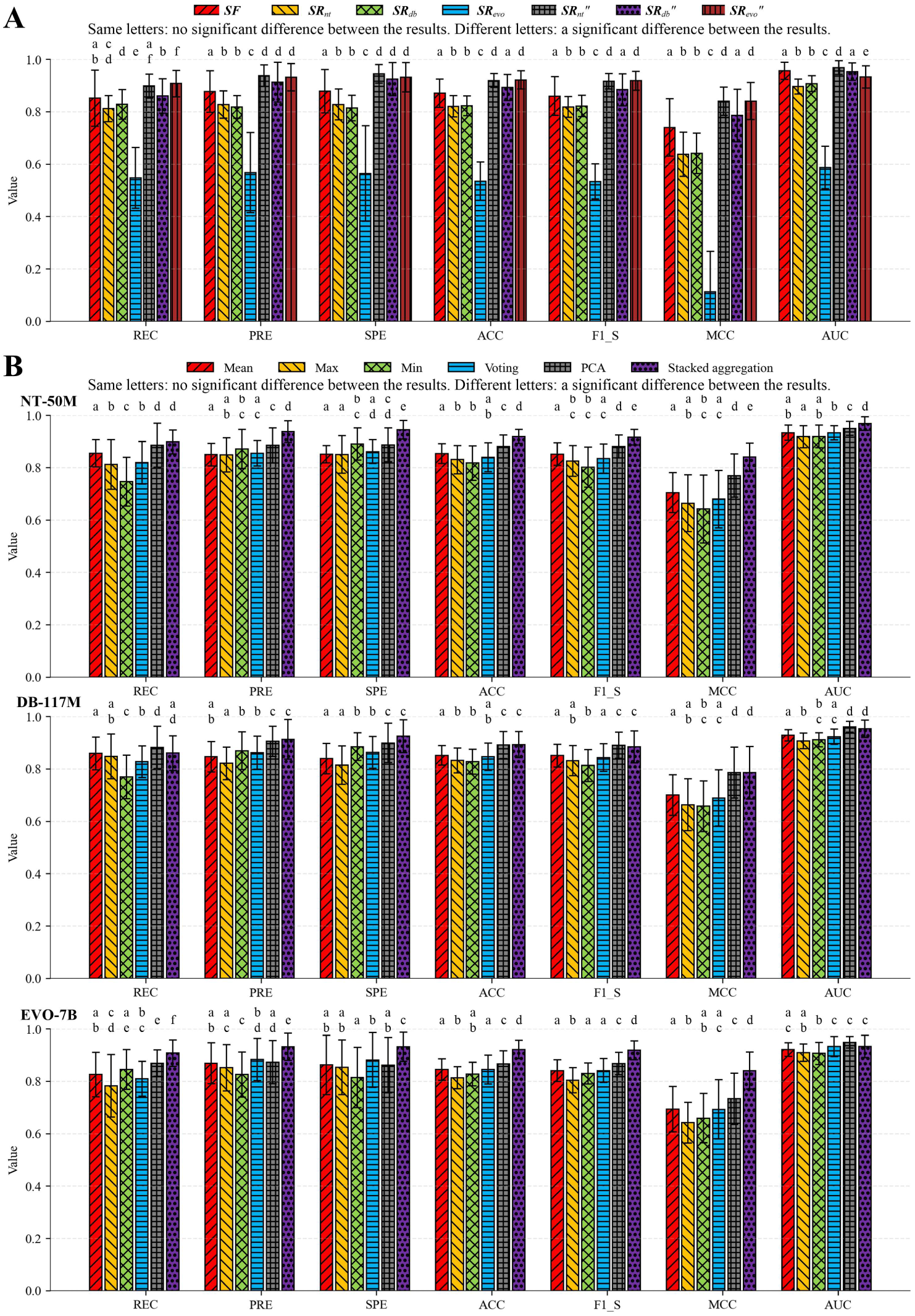
Effectiveness evaluation of enhanced representations and stacked aggregation classifier. The letters above the bars are results of Mann-Whitney U tests, where results labeled with the same letters indicate that there is no significant difference between them (*p*-value>0.05), and results labeled with different letters indicate that there is a significant difference between them (*p*-value≤0.05). (**A**) In the proposed method, fixing other modules and adopting different features/representations as inputs to the stacked aggregation classifier. ***SF*** denotes the engineered features. ***SR****_nt_*, ***SR****_db_* and ***SR****_evo_* denote the foundation model representations generated by Nucleotide Transformer-50M, DNABERT-2-117M and EVO-7B, respectively. ***SR****_nt_’’*, ***SR****_db_’’* and ***SR****_evo_’’* denote the enhanced representations, where the foundation model representations are generated by Nucleotide Transformer-50M, DNABERT-2-117M and EVO-7B, respectively. (**B**) In the proposed method, fixing other modules and employing different approaches based on the enhanced representation matrices for training and prediction. NT-50M denotes that the foundation model representations are generated by Nucleotide Transformer-50M. DB-117M denotes that the foundation model representations generated by DNABERT-2-117M. EVO-7B denotes the foundation model representations are generated by EVO-7B. Mean, Max, Min, and PCA denote the mean-taking approach, maximum-taking approach, minimum-taking approach, and PCA approach, respectively. Voting denotes the segment label voting approach. To ensure fairness, these approaches employ the LR model as classifier.

The classifier trained using engineered features obtains better results than those trained using foundation model representations. Classifiers trained using enhanced representations not only significantly outperform those trained using corresponding original foundation model representations but also surpass the classifier trained using engineered features in most cases. For the three foundation models, classifiers trained using enhanced representations derived from Nucleotide Transformer-50M and EVO-7B achieve superior performance compared to that trained using enhanced representations derived from DNABERT-2-117M. The stacked aggregation classifier obtains better results than classifiers based on other approaches overall, particularly when utilizing enhanced representations derived from Nucleotide Transformer-50M and EVO-7B. According to these results, we use the enhanced representations derived from Nucleotide Transformer-50M and EVO-7B for subsequent experiments.

### 3.3 Search for optimal hyperparameter set

We search the optimal hyperparameter set for the stacked aggregation classifier using the GridSearchCV method. This method systematically explores all specified hyperparameter combinations from predefined candidates, exhaustively training and evaluating each combination using cross-validation, and identifies the optimal hyperparameter set by selecting the combination that achieves the highest average score across all evaluations. We then configure the stacked aggregation classifier with the optimal hyperparameter set and retrain it using all samples from the training-validation set.

### 3.4 Comparison with other methods on balanced and imbalanced test sets

The proposed method is compared with iProbiotics [18], metaProbiotics [19] and MLC [48] on balanced and imbalanced test sets. IProbiotics and metaProbiotics provide the models trained using data from non-soil environments. MLC employs the GBRAP tool to extract the features from bacteria and archaea data to train a classifier. The proposed method only requires randomly selecting 100 segments per test sample for prediction, thus we perform 10 independent experiments to ensure reliability. For each experiment, segment group not entirely consistent with previous ones is selected per test sample to obtain the mean values for comparisons (Figure 5).

**Figure 5.**
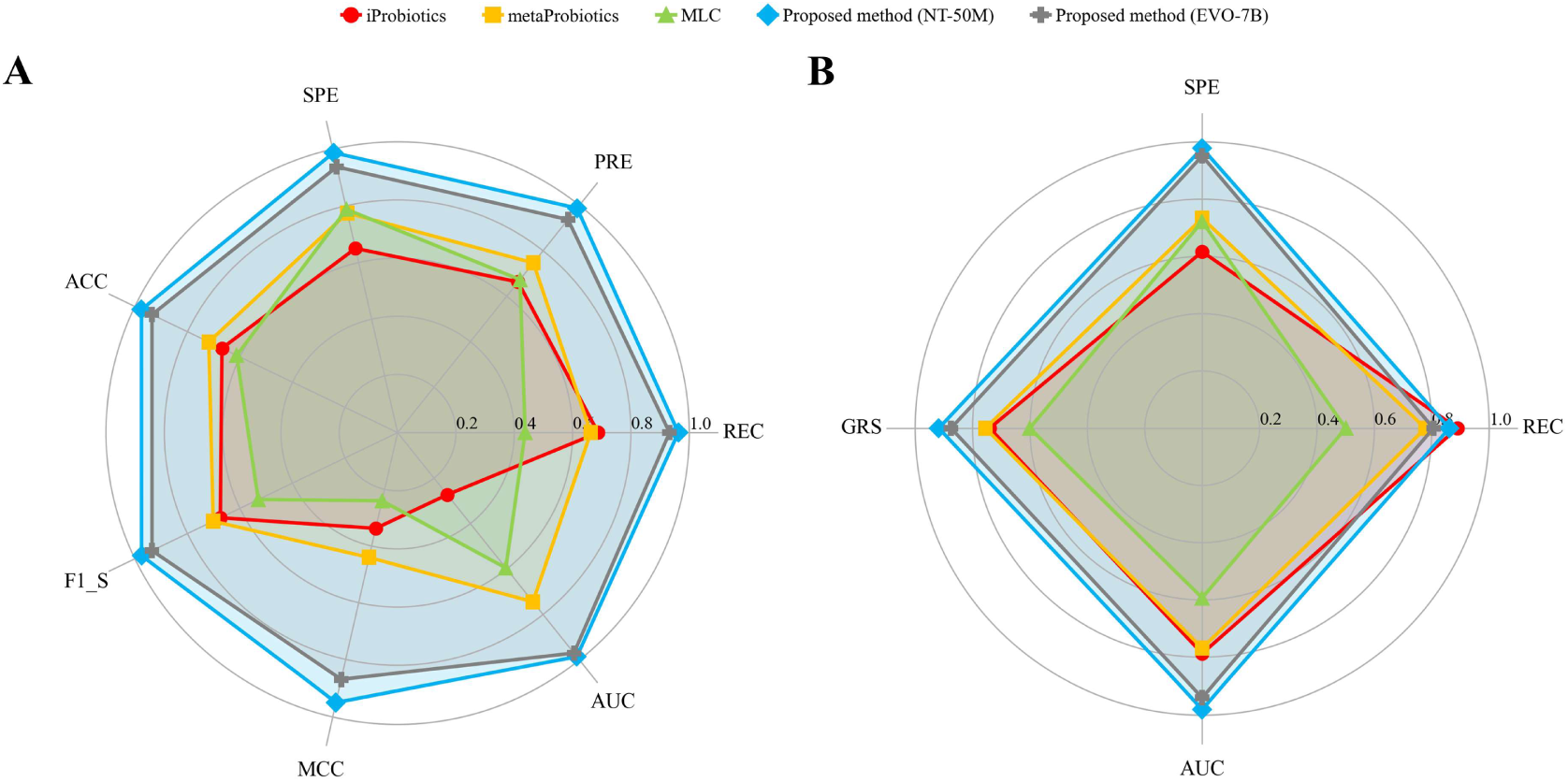
Comparisons of the proposed method with other methods. NT-50M and EVO-7B denote that in the proposed method, the foundation model representations are generated by Nucleotide Transformer-50M and EVO-7B, respectively. (**A**) Results on balanced test set. (**B**) Results on imbalanced test set.

On the balanced test set, the proposed method significantly outperforms other methods across all evaluation criteria. On the imbalanced test set, iProbiotics achieves the highest REC value, while the proposed method obtains best results in other evaluation criteria. Comparatively, utilizing enhanced representations derived from Nucleotide Transformer-50M in the proposed method enables better performance than utilizing those derived from EVO-7B.

### 3.5 Exploration of potential functional genes

We collate the labels and scores for all test samples predicted by the proposed method, and select two representative samples (GCA_001286965.1 and GCF_000242855.2) for gene function annotation and enrichment analysis. These samples are consistently predicted as probiotics across all independent experiments and are ranked within the top 25 by scores in each implementation of the proposed method (two implementations that the foundation model representations are derived from Nucleotide Transformer-50M and EVO-7B, respectively).

These two samples (*Bacillus amyloliquefaciens*) are reported to exhibit the capacity of solubilizing phosphate [49]. A total of 2,520 entries (involving 2,323 genes and 2,038 KO identifiers) and 2,532 entries (involving 2,333 genes and 2,049 KO identifiers) are effectively annotated for GCA_001286965.1 and GCF_000242855.2, respectively. Through mapping with functions, 8 genes in GCA_001286965.1 are assigned KO identifiers potentially associated with phosphate solubilization, such as K01113, K01077 and K01083, and the same cases are observed for GCF_000242855.2 (Table 2). Enrichment analysis results show that these genes may be involved in phosphate-solubilization-related pathways, such as map01100 (Metabolic pathways) and map02020 (Two-component system). These findings reveal genes that may play key roles in the capacity of solubilizing phosphate of *Bacillus amyloliquefaciens*.

**Table 2.**
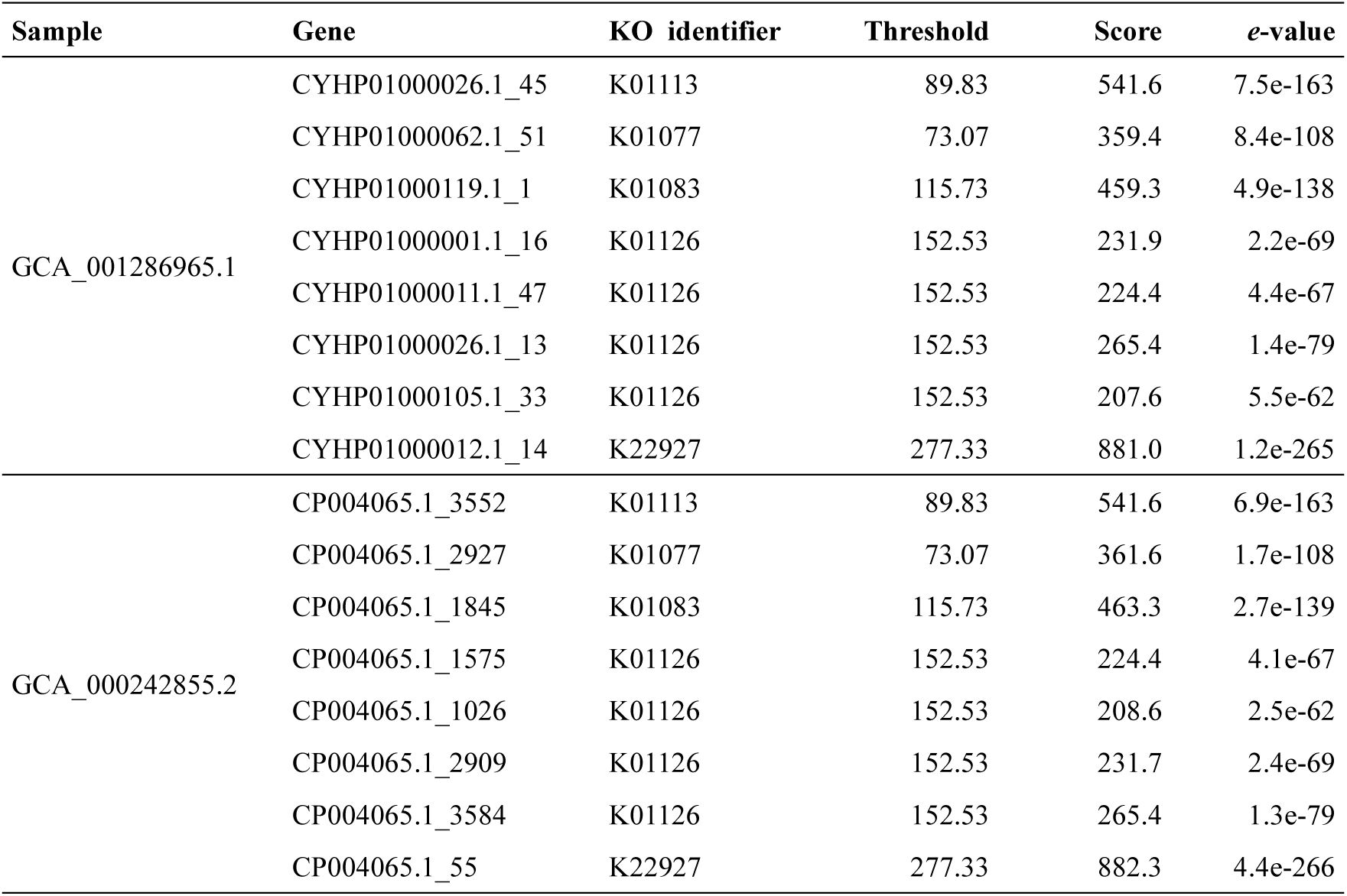
Function annotations of genes in GCA_001286965.1 and GCA_000242855.2 that the KO identifiers are potentially associated with phosphate solubilization.

| Sample | Gene | KO identifier | Threshold | Score | *e*-value |
| --- | --- | --- | --- | --- | --- |
| GCA_001286965.1 | CYHP01000026.1_45 | K01113 | 89.83 | 541.6 | 7.5e-163 |
|  | CYHP01000062.1_51 | K01077 | 73.07 | 359.4 | 8.4e-108 |
|  | CYHP01000119.1_1 | K01083 | 115.73 | 459.3 | 4.9e-138 |
|  | CYHP01000001.1_16 | K01126 | 152.53 | 231.9 | 2.2e-69 |
|  | CYHP01000011.1_47 | K01126 | 152.53 | 224.4 | 4.4e-67 |
|  | CYHP01000026.1_13 | K01126 | 152.53 | 265.4 | 1.4e-79 |
|  | CYHP01000105.1_33 | K01126 | 152.53 | 207.6 | 5.5e-62 |
|  | CYHP01000012.1_14 | K22927 | 277.33 | 881.0 | 1.2e-265 |
| GCA_000242855.2 | CP004065.1_3552 | K01113 | 89.83 | 541.6 | 6.9e-163 |
|  | CP004065.1_2927 | K01077 | 73.07 | 361.6 | 1.7e-108 |
|  | CP004065.1_1845 | K01083 | 115.73 | 463.3 | 2.7e-139 |
|  | CP004065.1_1575 | K01126 | 152.53 | 224.4 | 4.1e-67 |
|  | CP004065.1_1026 | K01126 | 152.53 | 208.6 | 2.5e-62 |
|  | CP004065.1_2909 | K01126 | 152.53 | 231.7 | 2.4e-69 |
|  | CP004065.1_3584 | K01126 | 152.53 | 265.4 | 1.3e-79 |
|  | CP004065.1_55 | K22927 | 277.33 | 882.3 | 4.4e-266 |

## 4. Discussions

Soil probiotics play indispensable roles in regulating agro-ecosystem functions, and reliable prediction of probiotics is crucial for discovering and cultivating novel functional strains. While current prediction methods predominantly focus on microorganisms in environments such as the human gut, specialized methods for soil probiotics remain conspicuously absent. This study presents a method based on foundation model representation enhancement and a stacked aggregation classifier to address this absence.

The classifier trained using engineered features demonstrate competitive performance, potentially due to their suitability for low-complexity classification tasks, and classifiers trained using enhanced representations achieve further breakthroughs. The stacked aggregation classifier employs a structure of two sub-classifiers to aggregate the enhanced representation vectors of multiple sequence segments into a single vector, significantly improving training effect compared to conventional approaches.

The results of iProbiotics, metaProbiotics and MLC verify the unreliability of models trained on non-soil microbial data for the prediction of soil probiotics. Gene function annotation and enrichment analysis for the predicted probiotics verify the strong support of the proposed method for biological discoveries. Collectively, the proposed method delivers robust performance for soil probiotic prediction. Its trained cross-attention module enables heterogeneous representation integration in related studies, and its stacked aggregation approach provides valuable references for long-sequence processing.

In future research, we will leverage the framework of the proposed method to conduct high-accuracy prediction for specific probiotic species, deeply investigate functional genes, and expect to derive new biological insights through wet-lab experiments.

## Key points

- Soil probiotics are predicted based on foundation model representation enhancement and stacked aggregation classifier.
- The foundation model representations are enhanced by deeply integrating domain-specific engineered features, enabling the training of a powerful classifier for a target task.
- A stacked aggregation classifier is designed that leverages partial sequence segments from a sample to predict its label, effectively processing long-sequences.
- The proposed method exhibits better prediction performance than other methods and provides strong support for biological discoveries.

## Data availability

All used data is public and downloaded from NCBI (https://www.ncbi.nlm.nih.gov/).

## Code availability

The proposed method is available at https://github.com/sunhaotong0605/SPP_FMRESAC.

